## Supplementary material for "A multi-channel electrophysiology approach to non-invasively and precisely record human spinal cord activity": SupplementaryMaterial.pdf

### *Analysis differences between preregistration and manuscript*

In this section, we list differences between the analyses proposed in the pre-registrations on OSF and the analyses we finally performed, as well as the reasons underlying these discrepancies.

Experiment 1: The preregistration stated that we aimed to also present SEPs at the channel with the strongest deflection. However, in the course of analyzing the data, we realized how well CCA was working on spinal data and decided that adding the time-course of the electrode with the strongest deflection would not bring additional value to the analysis, since CCA automatically incorporates the contribution of each channel to the SEP.

Experiment 1: The preregistration stated that we intended to investigate the relation between SEP amplitudes recorded at different levels of the somatosensory processing hierarchy. However, since we already show in Experiment 2 that there is mostly no such relation within one stimulation type, this analysis would not be very informative and we thus did not include it. Instead, we report a more informative analysis based on the data of Experiment 2, which allowed us to include different stimulation conditions (i.e., mixed, single-digit and double-digit stimulation)

Experiment 2: The preregistration stated that we wanted to include brainstem and Erb's point potentials in our analysis. However, due to low SNR we removed them from the results.

Experiment 2: The preregistration stated that we aimed to control for the difference in individual SEP latencies by taking the distance between the location of the recording and stimulation electrode into account. This was not necessary, because we used individual peak amplitudes and latencies in the present analysis.

Experiment 2: The preregistration stated that we intended to investigate the attenuation effect at the brainstem level (N14 and N30) as well. However, since the low SNR in the single-digit stimulation conditions did not allow for observing clear SEPs at the brainstem level, we were not able not to perform this analysis.

Experiment 2: The preregistration stated that for testing attenuation effects, we aimed to test the summed single-digit SEP-amplitudes against the double-digit amplitudes with a paired t-test. Since in the literature it is however more typical to calculate individual interaction ratios, we followed this approach and tested them against zero (Hsieh et al., 1995; Severens et al., 2010). However, for robustness we also performed paired t-tests and observed that this did not change the statistical decision (i.e., significant and non-significant comparisons remained in both analysis).

### *Mixed nerve results from Experiment 2*

We also aimed to replicate the main results from Experiment 1 (N=36) using the data from the mixed nerve conditions in Experiment 2 (N=24) and list these results in Supplementary Table 1.

### *Later spinal SEP components*

We aimed to replicate the late potentials observed in Experiment 1 with the data from the mixed nerve conditions in Experiment 2, using an identical approach. The following responses were identified via cluster-based permutation testing (after the early potentials, which are ignored here): i) in the hand mixed condition, we identified a cervical cluster directly after the N13 component between 19 ms and 24 ms ($p_{mcc} = 0.012$; channels: S3, S6, S7, S9, S11, S14, S18) that has higher activity during stimulation than during rest and ii) in the foot-mixed condition, we identified a positive cluster directly after the N22 component between 29 ms and 35 ms ($p_{mcc}$ = 0.004; channels: S22, S23, S26, L1, S28, S30, S32). This replicated the main results observed in Experiment 1, with the exception of the late potential displayed in Supplementary Figure 1.

### *Sensory nerve stimulation results*

Detailed results from the sensory nerve stimulation conditions in Experiment 2 are listed in Supplementary Table 2. Comparisons with mixed nerve stimulation revealed that peaks following double finger stimulation (fingers1&2) occurred 3.91 ms (peripheral), 4.30 ms (spinal), and 3.90 ms (cortical) later than those following hand-mixed stimulation and were 76% (peripheral), 66% (spinal), and 59% (cortical) smaller in amplitude. Potentials following double toe stimulation (toes1&2) occurred 6.05 ms (peripheral), 7.63 ms (spinal), and 8.39 ms (cortical) later compared to mixed nerve stimulation and were 88% (peripheral), 63% (spinal), and 43% (cortical) smaller in amplitude. Statistical comparisons of these differences are listed in Supplementary Table 3.

### *Shared response properties across the somatosensory hierarchy*

We here provide more extensive details regarding the investigation on whether response properties are shared across the somatosensory hierarchy.

Across the hand stimulation conditions, cortical SEP amplitudes were predicted by spinal SEP amplitudes, and spinal SEP amplitudes were predicted by peripheral NAP amplitudes ($\beta_{ESG}$ = 0.03, $t(145171.5) = 10.01$, $p < 0.001$ and $\beta_{periphery}$ = 0.02, $t(142032.7) = 8.40$, $p < 0.001$). Adding the factor *stimulation condition* to the models revealed that both these relationships were driven by effects of the type of stimulation on cortical SEP amplitudes ($\beta_{finger1}$ = 0.72, $t_{finger1}(145163.8)$ = 98.35, $\beta_{finger2}$ = 0.62, $t_{finger2}(145185.0)$ = 89.15, $\beta_{fingers1\&2}$ = 0.47, $t_{fingers1\&2}(145185.0)$ = 66.99, all $p < 0.001$), as well as on spinal SEP amplitudes ($\beta_{finger1}$ = 0.21, $t_{finger1}(136741.6)$ = 26.11, $\beta_{finger2}$ = 0.19, $t_{finger2}(141498.6)$ = 24.22, $\beta_{fingers1\&2}$ = 0.15, $t_{fingers1\&2}(141288.6)$ = 19.87, all $p < 0.001$); effect contrasts with reference level *mixed nerve stimulation*. At the same time, the effects of spinal SEP on cortical SEP amplitude and of peripheral NAP amplitude on spinal SEP amplitude were no longer significant and thus fully explained by *stimulation condition* ($\beta_{ESG}$ = 0.001, $t(145177.4)$ = 0.72, $p$ = 0.47 and $\beta_{periphery}$ = -0.00, $t(142364.2)$ = -0.42, $p$ = 0.672). Hence, *finger1*, *finger2*, as well as *fingers1&2* stimulations all resulted in differential amplitudes as compared to *mixed nerve* stimulation, both on the spinal as well as on the cortical level, and this amplitude variance was fully shared among the processing levels, explaining single-trial covariation across periphery, spinal cord and cortex.

A similar picture emerged for foot stimuli: cortical SEP amplitudes were predicted by spinal SEP amplitudes ($\beta_{ESG}$ = -0.04, $t(151307.0)$ = -14.38, $p < 0.001$) and spinal SEP amplitudes were predicted by peripheral NAP amplitudes ($\beta_{periphery}$ = 0.02, $t(151223.8)$ = 7.94, $p < 0.001$) when not controlling for stimulation conditions; please note that the negative sign of $\beta_{ESG}$ reflects the fact that spinal SEP amplitudes are measured as negative potentials while the first cortical SEP in the foot region, the P40, is a positive peak. When adding the factor *stimulation condition*, again, all types of stimulation affected the cortical level ($\beta_{toe1}$ = -0.61, $t_{toe\ 1}(151264.6)$ = -86.60, $\beta_{toe2}$ = -0.50, $t_{toe2}(151264.8)$ = -70.15, $\beta_{toes1\&2}$ = -0.32, $t_{toes1\&2}(151268.8)$ = -46.32, all $p < 0.001$) as well as the spinal level ($\beta_{toe1}$ = 0.33, $t_{toe\ 1}(140802.2)$ = 43.82, $\beta_{toe2}$ = 0.33, $t_{toe2}(140806.3)$ = 43.51, $\beta_{toes1\&2}$ =

.26, $t_{toes1\&2}$(141379.1) = 35.86, all $p < 0.001$). While the factor *stimulation condition* fully accounted for the effect of peripheral NAP on spinal amplitude, which was no longer existent ($\beta_{periphery} = 0.00$, $t$(151323.8) = 0.00, $p > 0.99$), a main – though slightly attenuated – effect of spinal amplitude on cortical amplitude still remained ($\beta_{ESG} = -0.02$, $t$(151320.4) = -3.72, $p < 0.001$). Additionally, small interaction effects on cortical amplitudes emerged between spinal amplitudes and toe2 stimulation, $\beta_{ESG * toe 2} = 0.02$, $t_{ESG * toe2}$(151314.1) = 3.02, $p_{ESG * toe2} = 0.003$, as well as between spinal amplitudes and toes 1 & 2 stimulation, $\beta_{ESG * toes1\&2} = 0.02$, $t_{ESG * toes1\&2}$(151314.5) = 2.99, $p_{ESG * toes1\&2} = 0.003$.

Taken together, the effects of different stimulation types (i.e., *mixed nerve, finger1/toe1*, *finger2/toe2*, *fingers1&2/toes1&2*) seem to propagate through the somatosensory processing hierarchy, jointly affecting the amplitudes of peripheral NAPs, spinal cord responses, and initial cortical potentials in the primary somatosensory cortex. This observation applied to both hand and foot stimulation, though with additional effects of spinal amplitudes on cortical amplitudes beyond the effect of stimulation condition in foot stimuli.

### *Placement of recording reference*

While other ESG studies have also used our choice of reference position for the recording of cervical and lumbar SEPs [1–9], the choice of reference position is always a compromise. For spinal recordings a non-cephalic reference as used here is generally suggested, but studies often use different references for cervical and lumbar recordings, such as the acromion for cervical or the pelvic bone for lumbar recordings. We wanted to use a reference position that is i) not lateralized, ii) ideal for both cervical and lumbar recordings and iii) positioned on a bone (not on muscle) and thus selected the spinous process of the 6th thoracic vertebra after running several pilot recordings with different reference positions.

**List of legends**

**Fig. S1.** Grand-average over all participants in the foot-mixed condition and in simulated epochs from rest data. The plotted signal is an average over all channels that are part of the identified cluster (channels displayed as red dots on the top left). The gray area between 126-132 ms identifies the time range in which the two signals are statistically different; note that this result did not replicate in Experiment 2.

**Table S1.** Group-level descriptive statistics for SEP- and NAP-amplitudes, latencies and SNR (mean and standard error of the mean) and one-sample t-test of SEP- and NAP-amplitudes in the hand-mixed and foot-mixed conditions of Experiment 2 (N = 24). Note that we only focused on the major peripheral, spinal and cortical components here for replication purposes and thus do not report Erb's point and brainstem potentials. Abbreviations: vr = ventral reference, tr = thoracic reference, CCA – canonical correlation analysis, SEP = somatosensory evoked potential, NAP = nerve action potential, # = number of participants in which potential was visible at the individual level, SNR = signal-to-noise ratio).

**Table S2.** Group-level descriptive statistics for SEP- and NAP-amplitudes, latencies and SNR (mean and standard error) and one-sample t-test of SEP- and NAP-amplitudes in all hand-sensory and foot-sensory conditions of Experiment 2 (vr = ventral reference, tr = thoracic reference, CCA = canonical correlation analysis, # = number of participants with potentials visible at the individual level).

**Table S3.** Paired t-test for the comparisons between hand-mixed and fingers1&2 conditions or foot-mixed and toes1&2 conditions. Tested were the amplitudes and the latencies of the SEPs and peripheral NAPs. Data come only from Experiment 2 (vr = ventral reference, tr = thoracic reference, CCA = canonical correlation analysis).

**S1 Data.** Each sheet (five sheets for Figure 2A, five sheets for Figure 2B) contains every participant's electrophysiological data (ENG, ESG, EEG), which together give rise to one grand-average trace reported in the figure. Each row reflects one timepoint and each column represents the amplitude of one participant's average evoked response.

**S2 Data. Sheets Figure_3A and Figure_3E.** Each sheet contains every participant's ESG data, which together give rise to one grand-average trace reported in the figure. Each row reflects one timepoint and each column represents the amplitude of one participant's average evoked response, first for single-channel data and then for CCA data. **Sheets Figure_3B and Figure_3F.** Each sheet contains every participant's ESG data, which together give rise to the grad-average isopotential plots. Each row corresponds to one channel and each column corresponds to one participant. **Sheets Figure_3C_sub-1-18, Figure_3C_sub-19-36, Sheets Figure_3G_sub-1-18, Figure_3G_sub-19-36.** Each sheets contains every participant's ESG data, which together give rise to one grand-average time-frequency plot. The rows correspond to frequencies and the columns correspond to time-points per participant. Please note that we had to create two sheets for each panel due to data-size limitations. **Sheets Figure_3D and Figure_3H.** Each sheet contains every participant's ESG data, which together give rise to one grand-average trace reported in the figure. Each row reflects one timepoint and each column represents the amplitude of one participant's average evoked response (average over all channels that are part of the identified cluster), first for stimulation and then for resting-state data. The last column depicts time-points of significant difference as established by the cluster-based permutation test.

**S3 Data. Sheets Figure_4A and Figure_4E.** Each sheet contains the ESG-SNR values of each participant (columns), with single channel data being represented in the upper row and CCA data being represented in the lower row. **Remaining six sheets.** Each sheet contains the individual participant ESG data underlying panels B-D and F-H (sub-006, sub-014, sub-021), with the first three sheets depicting cervical data and the last three sheets depicting lumbar data. Each row corresponds to one trial and each column corresponds to one time-point, first presented for the single-channel data and then for the CCA data.

**S4 Data.** Each sheet (one sheet for Figure 5A, one sheet for Figure 5B) contains every participant's ESG data, which together give rise to one grand-average trace reported in the figure. Each row reflects one timepoint and each column represents the amplitude of one participant's average evoked response after CCA, first for mixed and then for sensory nerve data.

**S5 Data.** Each sheet (three sheets for Figure 6A, three sheets for Figure 6B) contains every participant's electrophysiological data (ENG, ESG, EEG), which together give rise to one grand-average trace reported in the figure. Each row reflects one timepoint and each column represents the amplitude of one participant's average evoked response after CCA, first for digit 1 stimulation, then for digit 2 stimulation and finally for digit 1&2 stimulation.

**S6 Data. Sheets Figure_8A_Upper and Figure_8A_Lower.** Depicted are EEG data, with each row reflecting one timepoint and each column representing the amplitude of one participant's average evoked response from one electrode. **Sheet Figure_8B.** Depicted are ESG data, with each row reflecting one timepoint and each column representing the amplitude of one participant's average evoked response of component 1 after CCA for each given timepoint. **Sheet Figure_8C.** Depicted are ESG data, with each row reflecting one timepoint and each column representing the amplitude of one participant's evoked response based on the following subsets of trials in component 1 after CCA: _1: every fourth available trial beginning with trial 1; _2: every fourth available trial beginning with trial 2; _3: every fourth available trial beginning with trial 3; _4: every fourth available trial beginning with trial 4. **Sheet Figure_8D.** Depicted are ESG data, with the data being the same as those displayed in panel B, though in this case rather than plotting the average across all participants, each panel depicts a one participant's evoked response across all trials.

**S7 Data.** This sheet contains every participant's ESG data, which together give rise to one grand-average trace reported in the figure. Each row reflects one timepoint and each column represents the amplitude of one participant's average evoked response (average over all channels that are part of the identified cluster), first for stimulation and then for resting-state data. The last column depicts time-points of significant difference as established by the cluster-based permutation test.

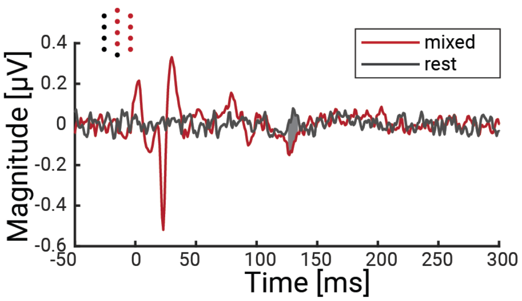


**Fig. S1.** Grand-average over all participants in the foot-mixed condition and in simulated epochs from rest data. The plotted signal is an average over all channels that are part of the identified cluster (channels displayed as red dots on the top left). The gray area between 126-132 ms identifies the time range in which the two signals are statistically different; note that this result did not replicate in Experiment 2**.** The data underlying this figure can be found in the Supplementary Material (S7 Data).

**Table S1.** Group-level descriptive statistics for SEP- and NAP-amplitudes, latencies and SNR (mean and standard error of the mean) and one-sample t-test of SEP- and NAP-amplitudes in the hand-mixed and foot-mixed conditions of Experiment 2 (N = 24). Note that we only focused on the major peripheral, spinal and cortical components here for replication purposes and thus do not report Erb's point and brainstem potentials. Abbreviations: vr = ventral reference, tr = thoracic reference, CCA – canonical correlation analysis, SEP = somatosensory evoked potential, NAP = nerve action potential, # = number of participants in which potential was visible at the individual level, SNR = signal-to-noise ratio).

| SEP / NAP | # | Latency [ms] | Amplitude [µV / a.u.] | SNR | tstat | P | 95%-CI | Cohen's d |
|---|---|---|---|---|---|---|---|---|
| *Mixed median nerve stimulation (hand-mixed)* | | | | | | | | |
| N6 | 24 | 6.46 ± 0.10 | -2.61 ± 0.29 | 36.52 ± 12.23 | -8.93 | <0.001 | [-3.21; -2.01] | -1.82 |
| N13 (tr) | 24 | 13.46 ± 0.20 | -0.86 ± 0.07 | 9.37 ± 1.51 | -12.31 | <0.001 | [-1.01; -0.72] | -2.51 |
| N13 (vr) | 24 | 13.75 ± 0.17 | -1.38 ± 0.09 | 14.36 ± 1.98 | -16.09 | <0.001 | [-1.55; -1.20] | -3.28 |
| N13 (CCA) | 24 | 13.58 ± 0.19 | -0.39 ± 0.04 | 24.01 ± 3.64 | -10.40 | <0.001 | [-0.46; -0.31] | -2.12 |
| N20 (CCA) | 24 | 19.79 ± 0.17 | -1.10 ± 0.08 | 24.07 ± 2.28 | -13.80 | <0.001 | [-1.26; -0.93] | -2.82 |
| *Mixed tibial nerve stimulation (foot-mixed)* | | | | | | | | |
| N8 | 22 | 9.54 ± 0.16 | -0.99 ± 0.16 | 13.30 ± 4.49 | -6.20 | <0.001 | [-1.32; -0.66] | -1.27 |
| N22 (tr) | 24 | 24.21 ± 0.36 | -0.57 ± 0.07 | 6.20 ± 1.07 | -8.38 | <0.001 | [-0.71; -0.43] | -1.71 |
| N22 (vr) | 24 | 24.71 ± 0.43 | -0.48 ± 0.06 | 10.09 ± 1.70 | -8.53 | <0.001 | [-0.59; -0.36] | -1.74 |
| N22 (CCA) | 24 | 24.25 ± 0.32 | -0.48 ± 0.05 | 24.97 ± 5.66 | -9.46 | <0.001 | [-0.58; -0.37] | -1.93 |
| P40 (CCA) | 24 | 40.92 ± 0.58 | 1.17 ± 0.09 | 27.93 ± 3.07 | 12.80 | <0.001 | [0.98; 1.36] | 2.62 |

**Table S2.** Group-level descriptive statistics for SEP- and NAP-amplitudes, latencies and SNR (mean and standard error) and one-sample t-test of SEP- and NAP-amplitudes in all hand-sensory and foot-sensory conditions of Experiment 2 (vr = ventral reference, tr = thoracic reference, CCA = canonical correlation analysis, # = number of participants with potentials visible at the individual level).

| SEP / NAP | # | Latency [ms] | Amplitude [µV / a.u.] | SNR | tstat | p | 95%-CI | Cohen's d |
|---|---|---|---|---|---|---|---|---|
| | | ***Sensory median nerve stimulation (hand-sensory)*** | | | | | | |
| | | *Index finger (finger1)* | | | | | | |
| N6 | 23 | 10.29 ± 0.15 | -0.34 ± 0.04 | 9.90 ± 1.82 | -8.32 | <0.001 | [-0.43; -0.26] | -1.7 |
| N13 (tr) | 21 | 18.13 ± 0.26 | -0.19 ± 0.04 | 2.36 ± 0.35 | -5.02 | <0.001 | [-0.26; -0.11] | -1.03 |
| N13 (vr) | 23 | 18.13 ± 0.24 | -0.25 ± 0.04 | 3.33 ± 0.59 | -6.38 | <0.001 | [-0.33; -0.17] | -1.30 |
| N13 (CCA) | 21 | 18.17 ± 0.20 | -0.09 ± 0.01 | 3.50 ± 0.68 | -5.71 | <0.001 | [-0.12; -0.05] | -1.16 |
| N20 (CCA) | 24 | 23.79 ± 0.23 | -0.34 ± 0.04 | 17.08 ± 3.73 | -9.16 | <0.001 | [-0.41; -0.26] | -1.87 |
| | | *Middle finger (finger2)* | | | | | | |
| N6 | 23 | 10.17 ± 0.14 | -0.41 ± 0.05 | 9.42 ± 1.56 | -8.91 | <0.001 | [-0.50; -0.31] | -1.82 |
| N13 (tr) | 23 | 17.92 ± 0.28 | -0.24 ± 0.06 | 2.28 ± 0.45 | -3.84 | 0.001 | [-0.38; -0.11] | -0.78 |
| N13 (vr) | 23 | 17.75 ± 0.25 | -0.42 ± 0.06 | 3.31 ± 0.43 | -7.35 | <0.001 | [-0.53; -0.30] | -1.50 |
| N13 (CCA) | 24 | 17.79 ± 0.29 | -0.12 ± 0.01 | 4.90 ± 1.24 | -8.60 | <0.001 | [-0.15; -0.09] | -1.76 |
| N20 (CCA) | 24 | 23.71 ± 0.26 | -0.41 ± 0.04 | 22.29 ± 6.40 | -9.59 | <0.001 | [-0.50; -0.32] | -1.96 |
| | | *Index and middle finger (fingers1&2)* | | | | | | |
| N6 | 23 | 10.13 ± 0.13 | -0.76 ± 0.08 | 11.34 ± 1.84 | -9.51 | <0.001 | [-0.93; -0.60] | -1.94 |
| N13 (tr) | 22 | 17.38 ± 0.27 | -0.39 ± 0.05 | 4.36 ± 1.52 | -8.56 | <0.001 | [-0.49; -0.30] | -1.75 |
| N13 (vr) | 22 | 17.58 ± 0.22 | -0.61 ± 0.07 | 6.37 ± 1.11 | -8.89 | <0.001 | [-0.75; -0.47] | -1.81 |
| N13 (CCA) | 24 | 17.58 ± 0.25 | -0.16 ± 0.02 | 6.76 ± 1.89 | -9.09 | <0.001 | [-0.20; -0.13] | -1.86 |
| N20 (CCA) | 24 | 23.71 ± 0.24 | -0.58 ± 0.06 | 42.14 ± 15.78 | -9.92 | <0.001 | [-0.70; -0.46] | -2.02 |
| | | ***Sensory tibial nerve stimulation (foot-sensory)*** | | | | | | |
| | | *First toe (toe1)* | | | | | | |
| N8 | 20 | 15.46 ± 0.28 | -0.11 ± 0.02 | 4.05 ± 0.82 | -6.33 | <0.001 | [-0.14; -0.07] | -1.29 |
| N22 (tr) | 24 | 31.21 ± 0.60 | -0.17 ± 0.03 | 2.39 ± 0.69 | -6.63 | <0.001 | [-0.23; -0.12] | -1.35 |
| N22 (vr) | 24 | 31.25 ± 0.60 | -0.10 ± 0.02 | 1.72 ± 0.26 | -4.51 | <0.001 | [-0.14; -0.05] | -0.92 |
| N22 (CCA) | 22 | 31.38 ± 0.52 | -0.10 ± 0.01 | 3.61 ± 0.60 | -7.03 | <0.001 | [-0.13; -0.07] | -1.44 |
| P40 (CCA) | 24 | 49.83 ± 0.71 | 0.53 ± 0.07 | 26.84 ± 10.81 | 7.13 | <0.001 | [0.38; 0.68] | 1.46 |
| | | *Second toe (toe2)* | | | | | | |
| N8 | 20 | 15.71 ± 0.29 | -0.10 ± 0.02 | 5.49 ± 1.99 | -6.32 | <0.001 | [-0.13; -0.07] | -1.29 |
| N22 (tr) | 23 | 31.25 ± 0.58 | -0.21 ± 0.04 | 2.15 ± 0.32 | -5.78 | <0.001 | [-0.28; -0.13] | -1.18 |
| N22 (vr) | 23 | 31.04 ± 0.62 | -0.08 ± 0.02 | 2.81 ± 0.73 | -3.13 | 0.004 | [-0.13; -0.03] | -0.64 |
| N22 (CCA) | 23 | 31.38 ± 0.49 | -0.10 ± 0.01 | 4.20 ± 0.61 | -8.64 | <0.001 | [-0.13; -0.08] | -1.76 |
| P40 (CCA) | 23 | 50.42 ± 0.75 | 0.62 ± 0.08 | 26.84 ± 5.50 | 7.43 | <0.001 | [0.44; 0.79] | 1.52 |
| | | *First and second toe (toes1&2)* | | | | | | |
| N8 | 19 | 15.33 ± 0.28 | -0.19 ± 0.03 | 9.29 ± 2.92 | -6.95 | <0.001 | [-0.25; -0.13] | -1.42 |
| N22 (tr) | 23 | 31.21 ± 0.60 | -0.22 ± 0.02 | 3.87 ± 0.76 | -9.38 | <0.001 | [-0.26; -0.17] | -1.91 |
| N22 (vr) | 23 | 31.00 ± 0.56 | -0.18 ± 0.03 | 3.40 ± 0.72 | -5.27 | <0.001 | [-0.25; -0.11] | -1.08 |
| N22 (CCA) | 23 | 31.38 ± 0.48 | -0.18 ± 0.02 | 7.59 ± 1.72 | -8.44 | <0.001 | [-0.22; -0.14] | -1.72 |
| P40 (CCA) | 23 | 49.25 ± 0.73 | 0.81 ± 0.10 | 26.72 ± 5.66 | 8.42 | <0.001 | [0.61; 1.01] | 1.72 |

**Table S3.** Paired t-test for the comparisons between hand-mixed and fingers1&2 conditions or foot-mixed and toes1&2 conditions. Tested were the amplitudes and the latencies of the SEPs and peripheral NAPs. Data come only from Experiment 2 (vr = ventral reference, tr = thoracic reference, CCA = canonical correlation analysis).

| SEP / NAP | tstat | p | 95%-CI | Cohen's d |
|---|---|---|---|---|
| *Amplitude: Hand-mixed – fingers1&2* | | | | |
| N6 | -6.73 | <0.001 | [-2.41; -1.28] | -1.37 |
| N13 (tr) | -5.38 | <0.001 | [-0.65; -0.29] | -1.10 |
| N13 (vr) | -7.42 | <0.001 | [-1.14; -0.64] | -1.52 |
| N13 (CCA) | -9.56 | <0.001 | [-0.27; -0.17] | -1.95 |
| N20 (CCA) | -10.32 | <0.001 | [-0.62; -0.41] | -2.11 |
| *Latency: Hand-mixed – fingers1&2* | | | | |
| N6 | -28.20 | <0.001 | [-3.94; -3.40] | -5.76 |
| N13 (tr) | -18.10 | <0.001 | [-4.36; -3.47] | -3.70 |
| N13 (vr) | -18.10 | <0.001 | [-4.36; -3.47] | -3.70 |
| N13 (CCA) | -21.01 | <0.001 | [-4.39; -3.61] | -4.29 |
| N20 (CCA) | -32.88 | <0.001 | [-4.16; -3.67] | -6.71 |
| *Amplitude: Foot-mixed – toes1&2* | | | | |
| N8 | -5.35 | 0.001 | [-1.09; -0.48] | -1.09 |
| N22 (tr) | -5.50 | <0.001 | [-0.49; -0.22] | -1.12 |
| N22 (vr) | -5.08 | <0.001 | [-0.46; -0.19] | -1.04 |
| N22 (CCA) | -7.18 | <0.001 | [-0.38; -0.21] | -1.47 |
| P40 (CCA) | 4.00 | 0.001 | [0.17; 0.55] | 0.82 |
| *Latency: Foot-mixed – toes1&2* | | | | |
| N8 | -24.46 | <0.001 | [-6.24; -5.26] | -4.99 |
| N22 (tr) | -18.86 | <0.001 | [-7.86; -6.31] | -3.85 |
| N22 (vr) | -18.86 | <0.001 | [-7.86; -6.31] | -3.85 |
| N22 (CCA) | -20.82 | <0.001 | [-7.83; -6.42] | -4.25 |
| P40 (CCA) | -18.56 | <0.001 | [-9.26; -7.40] | -3.79 |
